# A fast computational model for circulatory dynamics: Effects of left ventricle-aorta coupling

**DOI:** 10.1101/2022.11.06.515370

**Authors:** Michael J. Moulton, Timothy W. Secomb

## Abstract

The course of diseases such as hypertension, systolic heart failure and heart failure with a preserved ejection fraction are affected by interactions between the left ventricle (LV) and the vasculature. To study these interactions, a computationally efficient, biophysically based mathematical model for the circulatory system is presented. In a four-chamber model of the heart, the LV is represented by a previously described low-order, wall volume-preserving model that includes torsion and base-to-apex and circumferential wall shortening and lengthening, and the other chambers are represented using spherical geometries. Active and passive myocardial mechanics of all four chambers are included. The cardiac model is coupled with a wave-propagation model for the aorta and a closed lumped-parameter circulation model. Parameters for the normal heart and aorta are determined by fitting to experimental data. Changes in the timing and magnitude of pulse wave reflections by the aorta are demonstrated with changes in compliance and taper of the aorta as seen in aging (decreased compliance, increased diameter and length), and resulting effects on LV pressure-volume loops and LV fiber stress and sarcomere shortening are predicted. Effects of aging of the aorta combined with reduced LV contractile force (failing heart) are examined. In the failing heart, changes in aortic properties with aging affect stroke volume and sarcomere shortening without appreciable augmentation of aortic pressure, and the reflected pressure wave contributes an increased proportion of aortic pressure.

## 1. INTRODUCTION

The aging aorta is marked by changes in stiffness, diameter, length, taper and impedance (Phan et al. 2016; Hickson et al. 2010). These changes in aortic properties lead to more prominent pulse wave reflections that arrive at the aortic inlet earlier in systole and affect LV stroke volume and sarcomere shortening (Park et al. 2020). Such effects are important in diseases such as hypertension (Laurent and Boutouyrie 2020; Sweitzer et al. 2013), left ventricular hypertrophy (London and Guerin 1999), systolic heart failure (Weber and Chirinos 2018) and heart failure with a preserved ejection fraction (HFpEF) (Chirinos 2017a; Chirinos 2017b). The effects of changes in aortic properties and function on LV contraction are termed ventricular-vascular interaction (VVI) (Borlaug and Kass 2008). VVI can be assessed using echocardiography and heart catheterization, and is commonly quantified using the varying elastance framework described first by Suga and Sagawa (Suga 2003). The ratio of aortic elastance (*E_a_* = end systolic aortic pressure/volume) to maximum ventricular elastance (*E_es_* = end systolic LV pressure/end systolic volume at different afterloads) at end systole is a measure of the coupling between aorta and LV (Shoucri 1998). However, the aorta and the systemic circulation create two distinct types of afterload on the LV, termed “pulsatile” and “resistive” (Weber and Chirinos 2018), and use of *E_a_/E_es_* has been criticized because it neglects the effects of pulsatile impedance (Chirinos 2017a).

The reservoir function of the aorta was recognized in the eighteenth century (Hales 1733). The classic Windkessel model (Frank 1899) accounts for the decay of diastolic pressure, but not for the augmentation of the systolic pressure with increased aortic stiffness and impedance (Wang et al. 2011). The importance of the pulse wave augmentation and early wave reflections in hypertension was demonstrated in clinical studies and mathematically (Pagoulatou and Stergiopulos 2017; Heusinkveld et al. 2019; Womersley 1955). In recent decades, wave theories for the aorta have been further developed (Parker 2009; Wang and Parker 2004; Pagoulatou and Stergiopulos 2017; Mynard and Smolich 2014), including wave separation analysis, which allows forward and backward wave components of the pulse wave to be deduced from pressure and flow waveforms (Hughes et al. 2013).

A model of the aorta based on the one-dimensional (1-D) partial differential Euler equations for propagation of flow and pressure waves can predict the timing and magnitude of pulse wave reflections (Wang and Parker 2004). Several authors have developed wave propagation models based on the nonlinear Euler equations (Wang et al. 2011; Matthys et al. 2007; Mynard and Smolich 2014; Pagoulatou and Stergiopulos 2017). These models predict that wave reflection due to aortic branching and taper generates complex wave patterns at the aortic inlet (Matthys et al. 2007). Approaches in which such a wave propagation model is coupled to a geometrically detailed model of the LV (Shavik et al. 2018; Chen et al. 2016) are computationally challenging because a stiff set of coupled partial differential equations representing both the LV and the aorta must be solved simultaneously. In a recent study, the CircAdapt model (Arts et al. 2005) was coupled to a 1-D aortic wave model (Heusinkveld et al. 2019). The reflected pressure wave was shown to be increased in the simulated hypertensive aorta.

For systematic investigation of effects of changing aortic and LV properties on hemodynamic and cardiac parameters, computationally efficient approaches are desirable. In previous work, we developed a spatially resolved low-order model of the LV (Moulton et al. 2017). In the present study, this LV model is coupled to a 1-D aortic wave propagation model based on the linearized Euler equations. The resulting computations run faster than real time on a standard personal computer. This approach is used to examine the behavior of the coupled LV and aorta with simulated aging of the aorta, with normal LV contractility and with reduced contractility representative of the failing heart. Aortic properties (stiffness, wave speed, taper, length and diameter) are obtained by fitting to experimental data (Hickson et al. 2010) for patients of different age groups (young, middle-aged and old). Effects of impaired or failing LV contractility are simulated by reducing the assumed maximum contractile force generated in the myocardium.

## 2. METHODS

### 2.1 Overview of model

The model is illustrated schematically in Figure 1. Key elements are as follows. (i) The LV is represented by a previously developed low-order axisymmetric model (Moulton et al. 2017), in which long- and short-axis lengthening and shortening, together with torsion, are described by three time-dependent parameters. (ii) The other three cardiac chambers (right ventricle [RV], right atrium [RA] and left atrium [LA]) are described by spherical models, in which a single parameter specifies the deformation. (iii) Flows through the heart valves are described by a 0-D model (Mynard et al. 2012) based on the Bernoulli equation. (iv) Pulse wave propagation in the aorta is modeled by the linearized Euler equations for conservation of mass and momentum. In a preliminary step, these equations are solved numerically for a short impulse of flow at the entrance of the aorta. In the simulation of cardiac cycles, the flows and pressures in the aorta are then found by convolution with the impulse response functions, which is computationally fast (Oppenheim et al. 1999). (v) A closed-loop, lumped-parameter model is used for the systemic and pulmonary arterial and venous systems. (vi) The resulting system is represented by a set of 20 ordinary differential equations that are solved by standard techniques. An overview of the model components is presented in the following sections, and further details including the governing equations are given in the Supplementary Materials.

**Figure 1.**
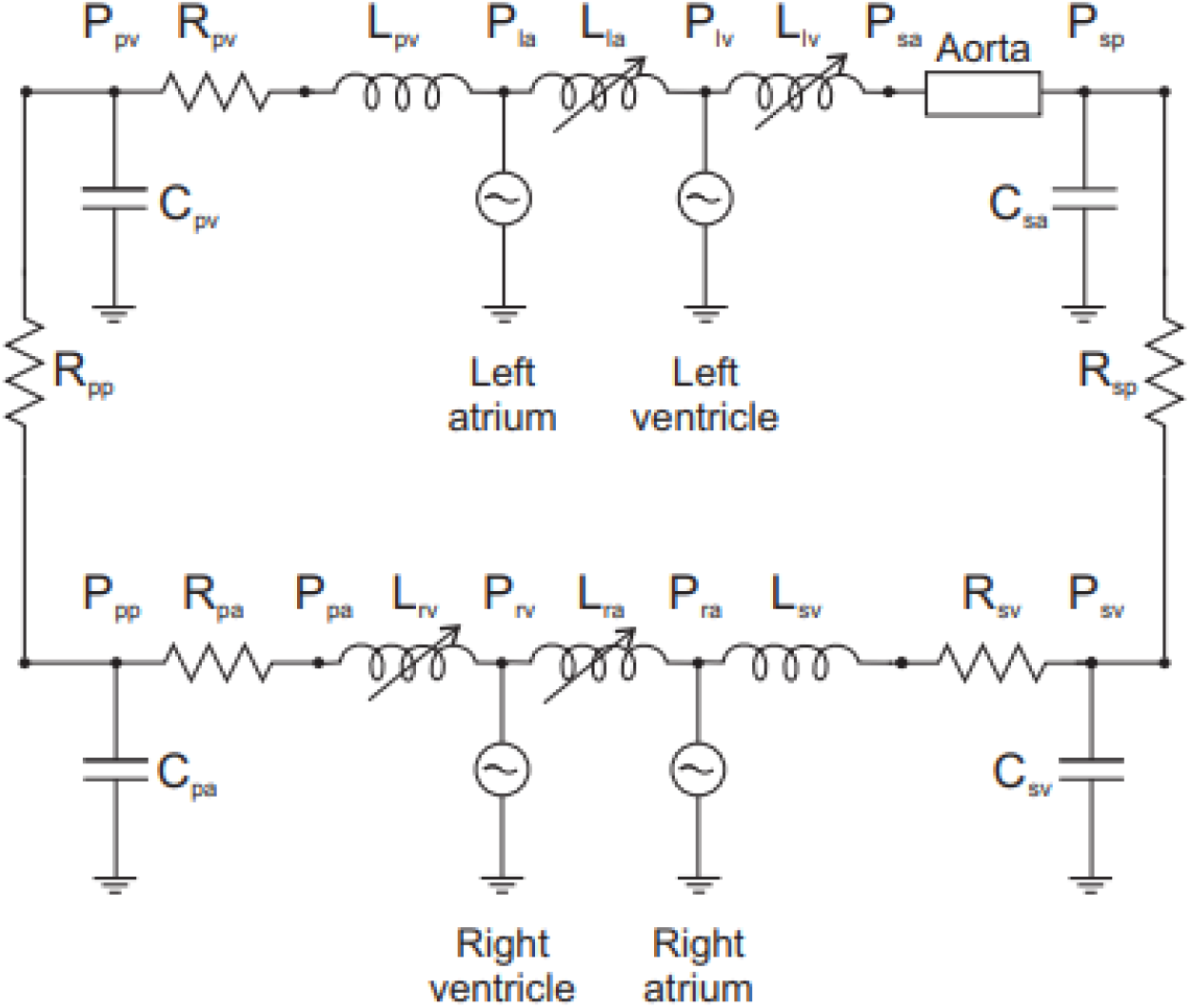
Schematic of circulatory system model, with components represented by corresponding electrical circuit elements. The four heart chambers are represented by thick-walled chambers with explicit representation of active and passive biophysical characteristics of myocardium. Valve models include variable inertial effects associated with flow, as indicated by the inductive elements *L_la_, L_lv_, L_ra_* and *L_rv_*. The aorta is represented by a 1-D wave propagation model. The other segments of the circulation are represented by lumped resistances, *R_sp_* and *R_sv_* for the systemic peripheral and venous vessels, and *R_pa_, R_pp_* and *R_pv_* for the pulmonary arterial, peripheral and venous vessels, and compliances (capacitances) *C_sa_* and *C_sv_* for the systemic arterial and venous vessels, and *C_pa_* and *C_pv_* for the pulmonary arterial and venous vessels. Inertial effects in the veins are indicated by inductive elements *L_sv_* and *L_pv_*.

### 2.2 Model for LV

The model for left ventricular dynamics has been described previously (Moulton et al. 2017). Briefly, the LV is represented as a thick-walled axisymmetric shell whose initial shape is a prolate spheroid (Figure 2A). Its deformation is fully specified by three time-dependent parameters (*a*_1_, *a*_2_, *a*_3_) that respectively describe base-to-apex, circumferential and torsional deformations using prolate spheroidal coordinates (*μ,v,ϕ*). The equation for the mapping from initial to deformed shapes is derived from the condition det(**F**) = *F_μμ_F_vv_F_ϕϕ_* =1 where *F* is the deformation gradient tensor, guaranteeing volume conserving deformation of the wall. A family of helical muscle fibers is introduced, with angles varying from endocardium to epicardium. Active force generation is aligned with the local fiber direction. The force varies with time according to an assumed activation function and with sarcomere length to account for the length-tension relationship (Figure 2B). Force-velocity dependence is introduced by including an activation-dependent viscous resistance to shortening. A viscoelastic model is used to describe the passive mechanical properties of the wall. The elastic component of stress assumes transversely isotropic properties with respect to the fiber direction, with an exponential type strain-energy function. The viscous stress is derived from the equations of viscous fluid motion in three dimensions. Force equilibrium is represented by the weak form of the equilibrium equations. This results in a set of three coupled linear ordinary differential equations in (*a*_1_, *a*_2_, *a*_3_) as functions of time.

**Figure 2.**
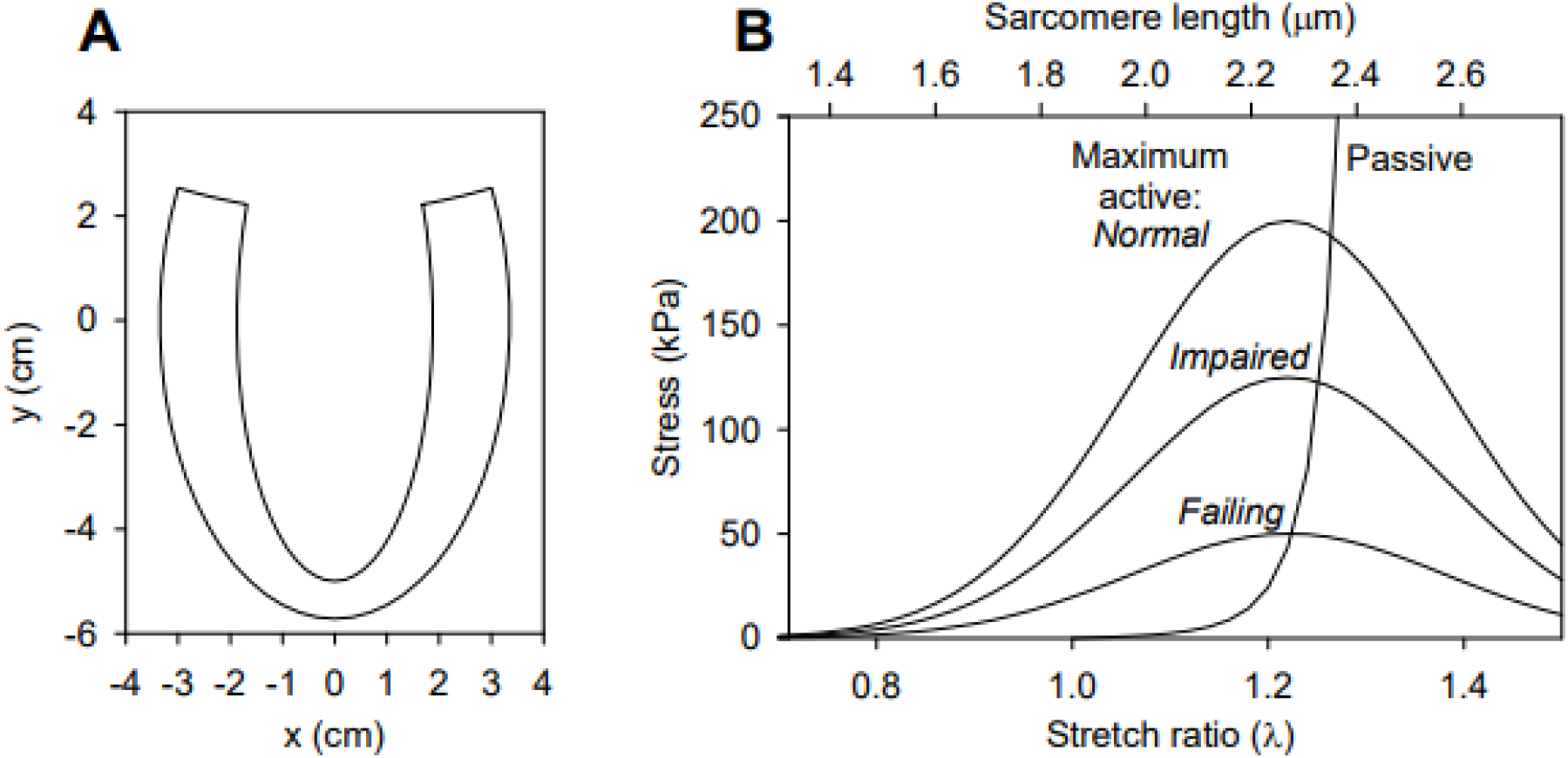
**A**. Profile of assumed axisymmetric LV shape in the initial configuration. The inner and outer surfaces are lines of constant *μ* in prolate spheroidal coordinates, and the upper boundary is a line of constant *v*. **B**. Assumed active and passive length-tension properties of cardiac muscle fibers. Results are given in terms of both sarcomere length and stretch ratio relative to initial configuration. Three different levels of active force generation are considered, as indicated.

### 2.3 Model for other cardiac chambers

The RV, LA and RA are each represented by a thick-walled sphere (LA and RA) or part of a sphere (RV), whose radius changes with time described by a parameter *a_j_, j* = 4,5,6, and with volume preserving wall deformations. The model is analogous to the one-fiber model (Arts et al. 1991; Arts et al. 2005), in that it explicitly represents the biophysical properties of muscle fibers within a simplified geometry. The force of fiber contraction is assumed to act isotropically in the surface of the sphere. As in the LV model, the fiber stress varies with time according to a prescribed activation function, and with sarcomere length according to the length-tension relationship, and force-velocity characteristics are represented by including a viscous resistance to shortening. Viscoelastic passive myocardial properties are again assumed. The weak form of the equations of mechanical equilibrium give rise to a single differential equation for *a_j_*(*t*).

### 2.4 Valve model

The four cardiac valves are represented by a 0-D Bernoulli-type model (Mynard et al. 2012; Arts et al. 2005), which can simulate normal and pathologic valve functions and reproduce flow and pressure curves observed in echocardiograms and invasive pressure measurements. Flow *q* through each valve is governed by an equation of the form

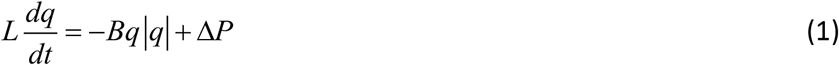

where Δ*P* is the pressure drop across the valve, *L* represents the inertia of the blood in the valve and the coefficient *B* relates the kinetic energy of blood passing through the valve to the square of the flow rate. The quantities *L* and *B* depend on the instantaneous effective valve area *A*, which varies between minimum and maximum values *A_closed_* and *A_open_* according to a function *ζ*(*t*) that represents the degree of valve opening:

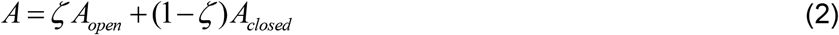

where 0 ≤ *ζ* ≤ 1. This function satisfies a dynamic equation

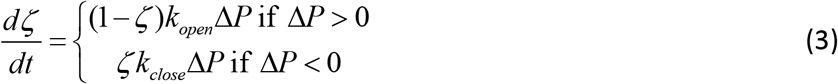

where *k_open_* and *k_close_* are rate constants of valve opening and closing. This model allows for a slight time lag between reversal of pressure drop and valve closing, and avoids discontinuities that occur in simpler models with instantaneous valve closure.

### 2.5 Spatially resolved aorta model

The diameter and wall properties of the aorta vary along its length, and several major arteries branch off. Changes in aortic diameter, gradients in aortic wall stiffness and branch vessels contribute to mismatching of wave impedance, causing wave reflection (Milnor 1975; Nichols et al. 1977; Alastruey et al. 2009; Secomb 2016). Although arterial branch points have previously been considered as the main source of wave reflections in the aorta (Papageorgiou and Jones 1987), reflections are generated continuously along the length of the aorta (Segers and Verdonck 2000; Milnor 1989). The reflected waves substantially affect the LV afterload (Segers and Verdonck 2000). In the present model, such reflections are represented by assuming that the cross-section area decays exponentially along the aorta.

A wave propagation model is required to represent these effects. In lumped-parameter models of the circulatory system, the Windkessel model is commonly used to represent the impedance of the aorta. With suitably tuned parameters, such models can predict approximate aortic pressure waveforms, but they do not represent the time delays associated with reflected waves. More realistic models for wave propagation in the aorta have been developed, including the effects of non-uniformity, branching and nonlinear elastic properties of vessels (Stergiopulos et al. 1993). Such detailed models generally require extensive computations.

In order to meet the goal of fast computational speed, a novel technique was developed based on impulse response functions for the Euler equations describing arterial wave propagation. The pressure *p*(*x, t*) and flow rate *q*(*x, t*) satisfy the equations for conservation of volume and momentum, linearized about a state where the flow is zero and the cross-section area is *A*_0_:

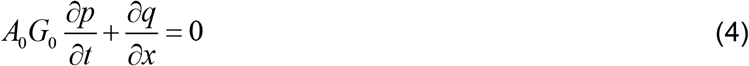

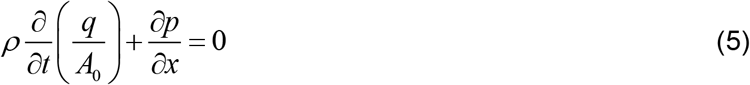

where *ρ* is the density of blood and *G*_0_ is the compliance of the aorta. The aorta is assumed to be tapered according to *A*_0_(*x*) = *A*_00_ exp(-*A*_01_ *x*), where *A*_00_ is the area at *x* = 0 and *A*_01_ defines the taper. Because the system is linear, the response to an arbitrary inflow waveform can be computed from the response to a brief impulse of flow. An inflow *q_in_*(*t*) lasting 0.02 s is imposed at *x* = 0 and a matched impedance condition is imposed at the downstream end (*x* = *L*). The equations are solved using a fourth-order Runge-Kutta scheme with a time step of 10^-4^ s and a grid spacing of 0.002*L*. Two impulse response functions are computed and stored: the reflected pressure *f_refl_*(*t*) at *x* = 0 and the transmitted flow *f_trans_*(*t*) at *x* = *L*. To obtain *f_refl_*(*t*), the pressure spike resulting from the imposed flow pulse is subtracted from *p*(0, *t*). The time-average is then subtracted, so that the resulting function has zero mean.

In the simulation of cardiac cycles, the flow rate entering the aorta is continuously convoluted with these impulse response functions, to obtain the reflected pressure wave and the aortic outflow rate. The steady component of the pressure at *x* = 0 is unaffected by wave reflection and depends on the peripheral resistance, so it is evaluated separately and added to the reflected wave, which has zero mean because *f_refl_*(*t*) has zero mean. In this way, the pulsatile and resistive loads (Weber and Chirinos 2018) are represented mathematically. An advantage of this method is that the reflected pressure wave is explicitly computed, so that its contribution to aortic pressure can be readily quantified.

### 2.6 Lumped-parameter circulatory system model

A lumped-parameter closed-loop model for the vascular system is used. The closed-loop system has the important advantage that interactions between all system components are explicitly represented. For example, changes in LV stroke volume affect flow rates and compartment volumes throughout the entire system.

The configuration is shown in Figure 1, using electrical circuit symbols to represent resistances, capacitances (compliances) and inductances (inertial effects). Five resistance elements are introduced, to represent the systemic peripheral vessels, systemic veins, pulmonary arteries, pulmonary peripheral vessels and pulmonary veins. Four compliance elements are introduced, to represent the systemic arteries (excluding the aorta), the systemic veins, the pulmonary arteries and the pulmonary veins. In addition, two elements, analogous to electrical inductances, are included to represent the inertia of blood in the systemic and pulmonary veins. This inertia is significant for limiting backflow in the veins during atrial contraction. The number of elements in the model is chosen to be sufficient to represent a closed loop and to provide an adequate representation of the preload and afterload on the heart chambers, while avoiding the complexity of more elaborate models. For each compliance element, flow conservation leads to a differential equation for the corresponding nodal pressure, and for each inertial element a differential equation governs the corresponding flow rate.

### 2.7 Solution method

The force balance equations for the LV, RV, LA, RA, together with the equations for valves and the lumped parameter elements make up a system of twenty ordinary differential equations. The reflected aortic pressure and aortic outflow rate, computed by convolution, are incorporated into this system of equations. The state variables include six for kinematics of the four chambers (*a*_1_ to *a*_6_), six for flow rates in and out of the four chambers, four for pressures at the compliant elements of the lumped model, and four for valve states. The solution is integrated using a second-order Runge-Kutta scheme, with a time step of 0.001 s. Simulation of 10 cardiac cycles takes less than 2 s on a personal computer (Dell XPS 9710 i9 Processor 2.50GHz; 64GB RAM).

### 2.8 Model parameters

Table 1 gives model parameters describing the heart chambers and the lumped-parameter circulatory model. Geometric parameters defining the reference shape for the LV (*a*_0_, *μ*_*in*0_, *μ*_*out*0_ and *v_up_*) and for the RA, LA and RV (*r*_*in*0_, *r*_*out*0_) are derived from a fit to echocardiographic images of a human volunteer (Moulton and Secomb 2014). The RV is represented by a fraction *f_RV_* of a sphere, to enable approximate matching of both the volume and the curvature of the chamber wall. Time constants (*T_c_, T_a_, T_ca_, T_cas_*) as defined in Table 1 are typical for a heart rate of 60 bpm. Parameters (*k_mLV_, k_mRV_, k_mLA_, k_mRA_*) that define the maximum active fiber stress in the four chambers were obtained by fitting to experimental data (ter Keurs et al. 2008). The active length-tension relationship is represented by a Gaussian function of sarcomere length. Effects of varying cardiac contractility are simulated by reducing *k_mLV_* to 62.5% and 25% of its *normal* value, corresponding to *impaired* and *failing* LV function. Parameters (*L*_*s0*_, *L*_*s*max_, *L_sw_*) that define the slack length, length at maximum force and width of the Gaussian, were estimated from the same data (ter Keurs et al. 2008). The force-velocity characteristics are represented by including an activation-dependent viscous resistance to contraction. The corresponding parameters (*k_vLV_, k_vRV_, k_vLA_, k_vRA_*) were obtained by fitting to data (de Tombe and ter Keurs 2012). Passive exponential, transversely isotropic stiffness parameters of the LV (*C_LV_, b_ff_, b_fx_, b_xx_*) were based on human data (Nordbø et al. 2014; Nordsletten et al. 2011). Passive exponential parameters that define the transversely isotropic stiffness of the RV and atria (*C_sph_, b_j_, b*_⊥_) were obtained from published experimental data (Schwartzman et al. 2013). The viscosity of the myocardial matrix *k_v_* was estimated previously (Moulton et al. 2017). Fiber angles ranged from −80° at the endocardium to +60° at the epicardium (Streeter and Hanna 1973). The resistances in the lumped-parameter circulatory model were estimated based on pressures typically observed in the various compartments and a resting cardiac output of 4.72 l/min. The assumed pulmonary artery compliance *C_PA_* is at the upper end of the published range (Thenappan et al. 2016). A similar value is assumed for the pulmonary venous compliance. The assumed systemic artery compliance *C_SA_*, when combined with the aortic compliance, is close to published values (Wohlfahrt et al. 2015; Thenappan et al. 2016). A large value is chosen for the systemic vein compliance *C_SV_*. The results are generally insensitive to the assumed venous compliances. Vein inertia parameters are estimated based on typical vein dimensions.

**Table 1.**
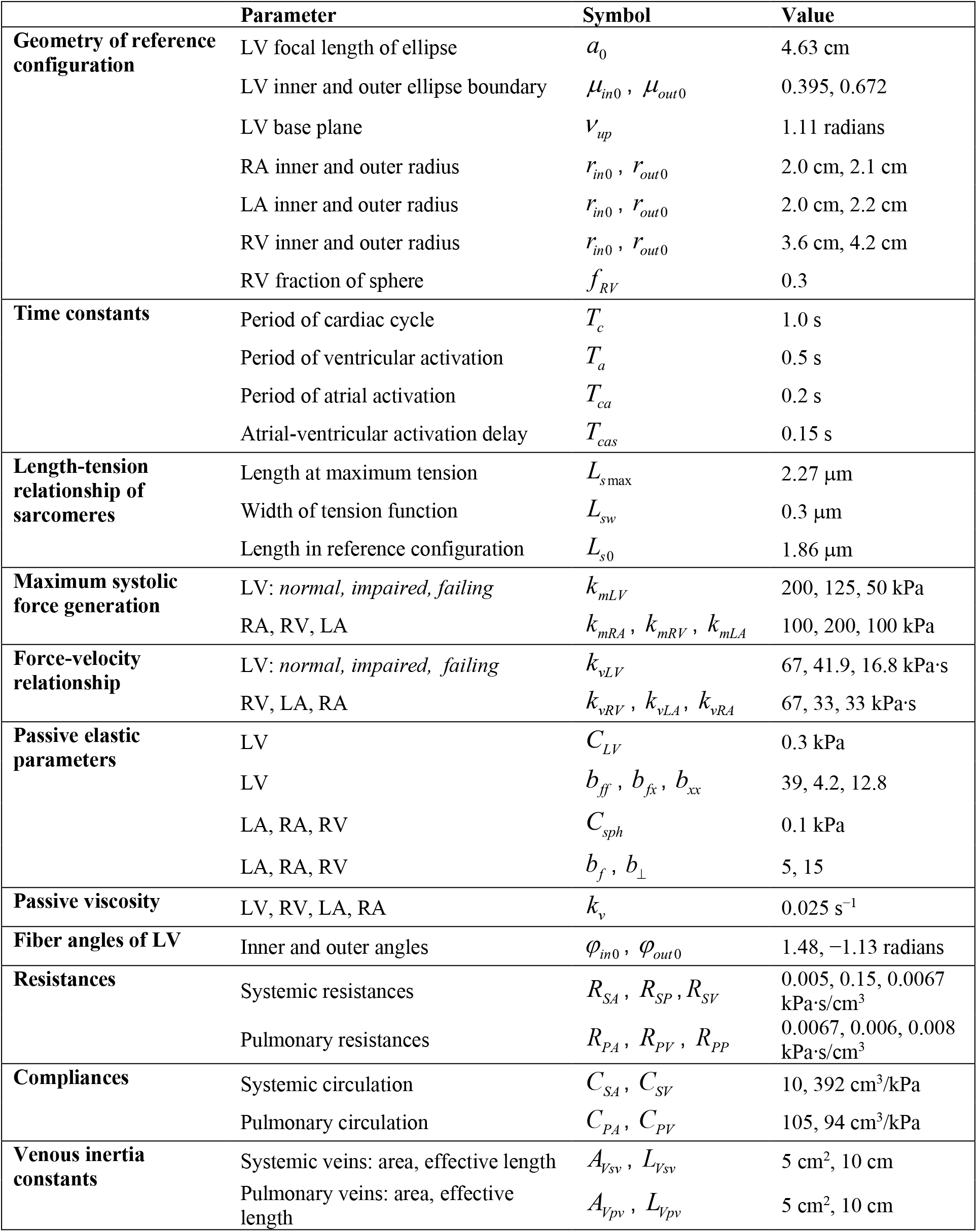
Model parameters: heart chambers and lumped-parameter model.

Table 2 gives model parameters describing the valves and the aorta. The maximal opening areas for each valve (open values of *A_aov_, A_mv_ A_pv_ A_tcv_*) were obtained from echocardiographic imaging data (Singh and Mohan 1994). Valve effective lengths (*V_Laov_, V_Lmv_ V_Lpv_ V_Ltcv_*) and kinetic time constants (opening and closing values for each valve, e.g., *K_aov,open_* were obtained from previous studies (Mynard et al. 2012; Arts et al. 2005). Aortic properties (length, compliance, inlet area and taper) are represented by parameters *L_aorta_, G, A*_00_ and *A*_01_, respectively. To simulate the aging aorta, data on pulse wave velocity, diameter, length and taper (regional change in diameter) at four locations along the aorta in 162 subjects (Hickson et al. 2010) were utilized to estimate *L_aorta_, G, A*_00_, *A*_01_ for *young, middle-aged* and *old* cohorts.

**Table 2.**
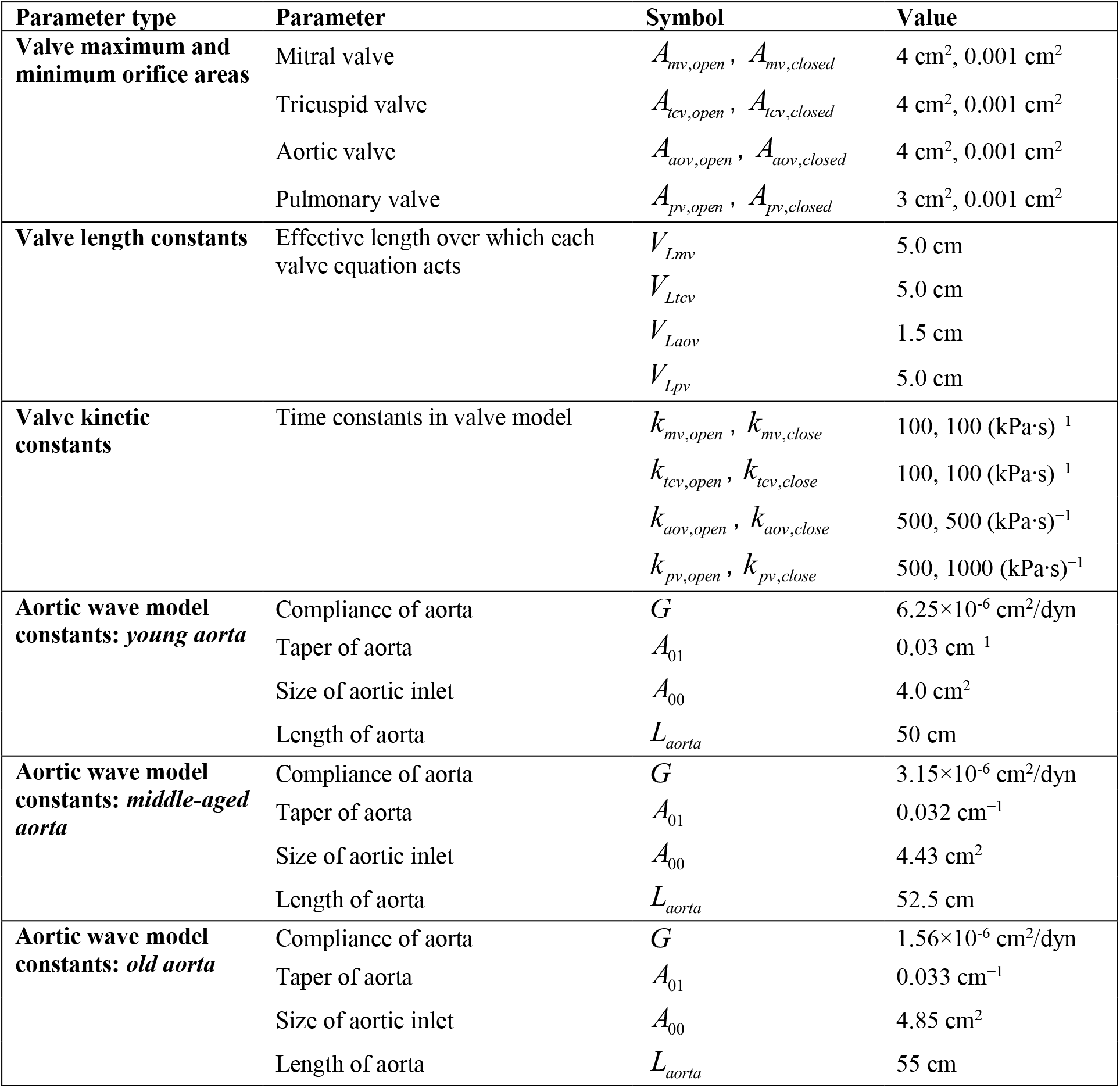
Model parameters: valves and aorta model.

## 3. RESULTS

### 3.1 PV loops

Figure 3 shows predicted PV loops in the LV for three conditions corresponding to *young*, *middle-aged* and *old* aortas. With increasing age, stroke volume is reduced, from 65.5 cm^3^ to 59.0 cm^3^, and pressure rises to an increasing peak in late systole. A right shift of the PV loop indicates increased preload pressures. In these results, all model parameters other than those describing the aorta were held constant.

**Figure 3.**
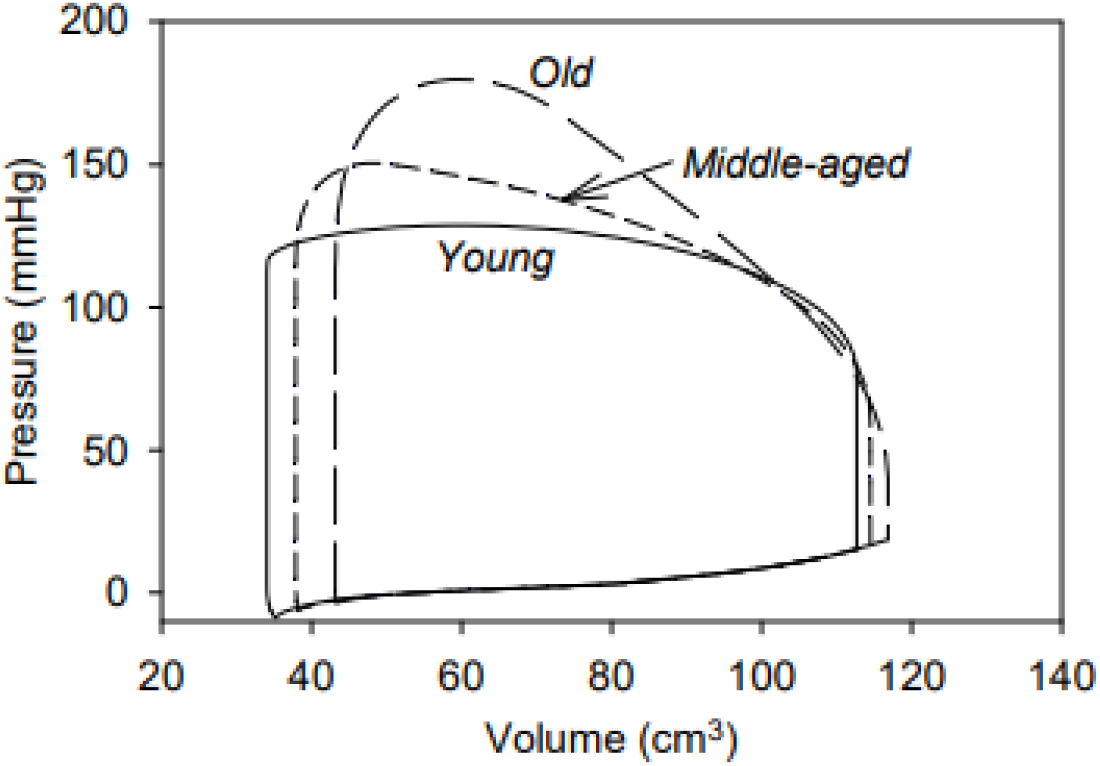
Predicted pressure-volume loops for the three sets of aortic properties, corresponding to *young, middle-aged*, and *old* humans, as defined in Table 2. Cardiac parameters are the same for each case.

### 3.2 Reflected wave properties

In Figure 4, effects of changes in aortic properties with age are further examined. Figure 4A shows the computed reflected pressure wave *f_refl_*(*t*) at the inlet of the aorta after a brief flow pulse at *t* = 0. With increasing aortic age and stiffness, the wave speed *c* = (*ρG*_0_)^-1/2^ in the aorta increases from 390 cm/s (*young*) to 550 cm/s (*middle-aged*) and 781 cm/s (*old*), resulting in the earlier arrival of the peak in the reflected wave. Furthermore, the increasing impedance of the aorta results in increased amplitude of the reflected wave. In simulations of the cardiac cycle, the function *f_refl_*(*t*) is convoluted with the inflow rate to the aorta to obtain the reflected pressure wave at the aortic inlet. Figure 4B shows the computed total pressure at the aortic inlet, together with the reflected component. With increasing aortic age, the reflected component reaches a higher peak, earlier in systole. The increase in total pressure closely parallels the increase in the reflected pressure wave. The diastolic decay of pressure is faster with aortic aging, because of the reduction in compliance, resulting in increased pulse pressure. With aging, the flow pulse at the aortic inlet shown in Figure 4C has reduced peak and prolonged duration. The flow pulse reaching the distal end of the aorta is shown in Figure 4D. With aging, the flow pulse arrives sooner due to increased wave speed, but is blunted in mid-systole due to the increased impedance of the aorta, resulting in prolongation of the flow pulse.

**Figure 4.**
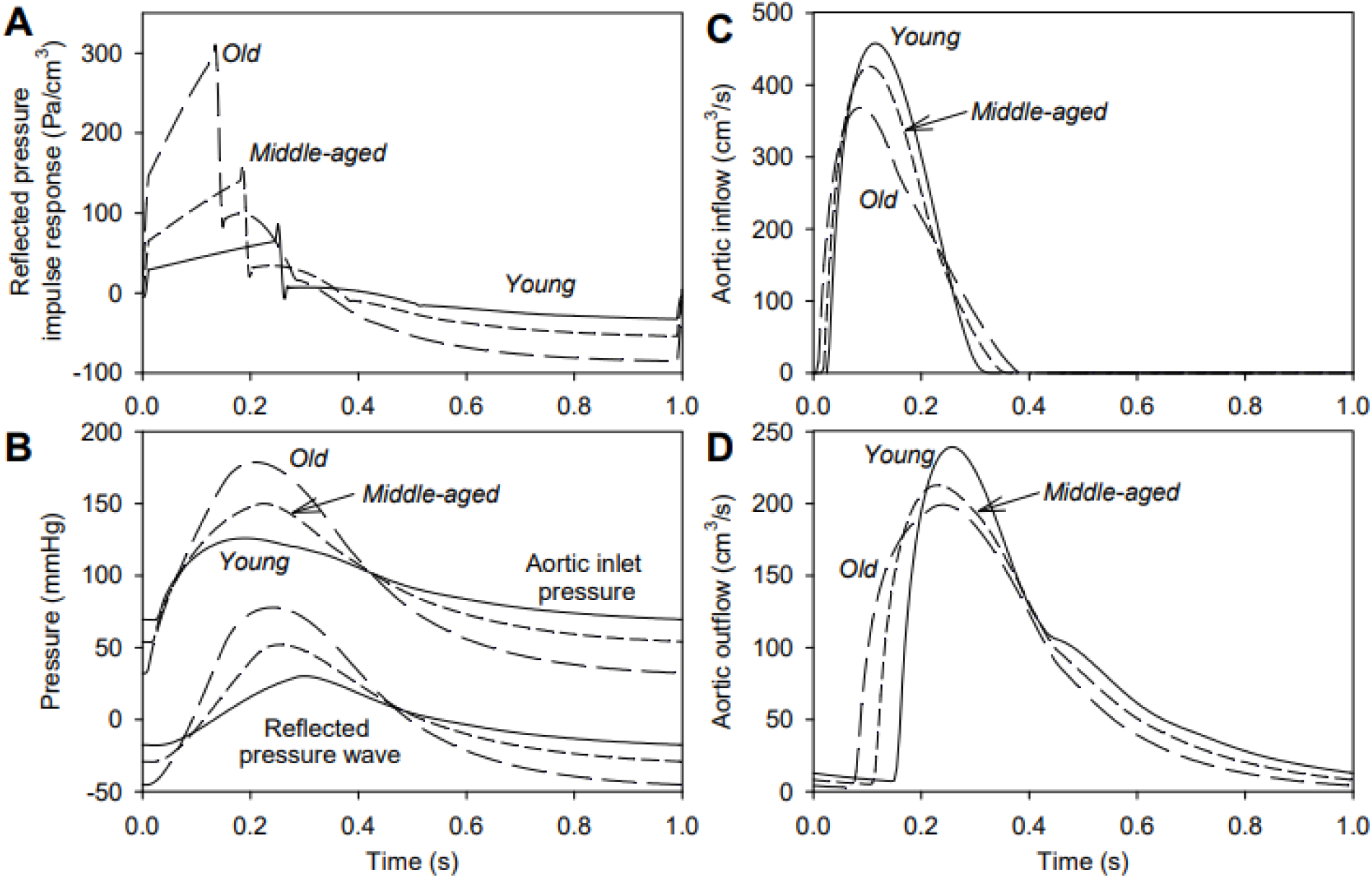
Predicted aortic pressures and flows for three sets of aortic properties (*young, middle-aged* and *old*). **A**. Reflected pressure at aortic inlet due to a brief flow pulse at t = 0. **B**. Actual pressure at aortic root, together with reflected component. **C**. Flow at the aortic inlet. **D**. Outflow from distal end of aorta.

### 3.3 Effects of aging aorta on sarcomeres

Figure 5A shows the active fiber stress, averaged over the LV wall volume, taking into account the force-velocity relationship. Figure 5B shows the corresponding variations in sarcomere length. With aging, the active fiber stress is increased. Two factors contribute to this effect. Firstly, the reduced rate of shortening in the older aorta results in increased force, according to the force-velocity relation. Secondly, the increased sarcomere length as systole progresses results in increased force generation because the sarcomeres are operating at a higher point on the length-tension curve (Figure 2B). With aging, the fiber stress reaches its peak later. As a consequence of the reduced rate of sarcomere shortening, overall shortening is reduced despite starting from a point of higher preload. The rate of relaxation of the sarcomeres is not affected.

**Figure 5.**
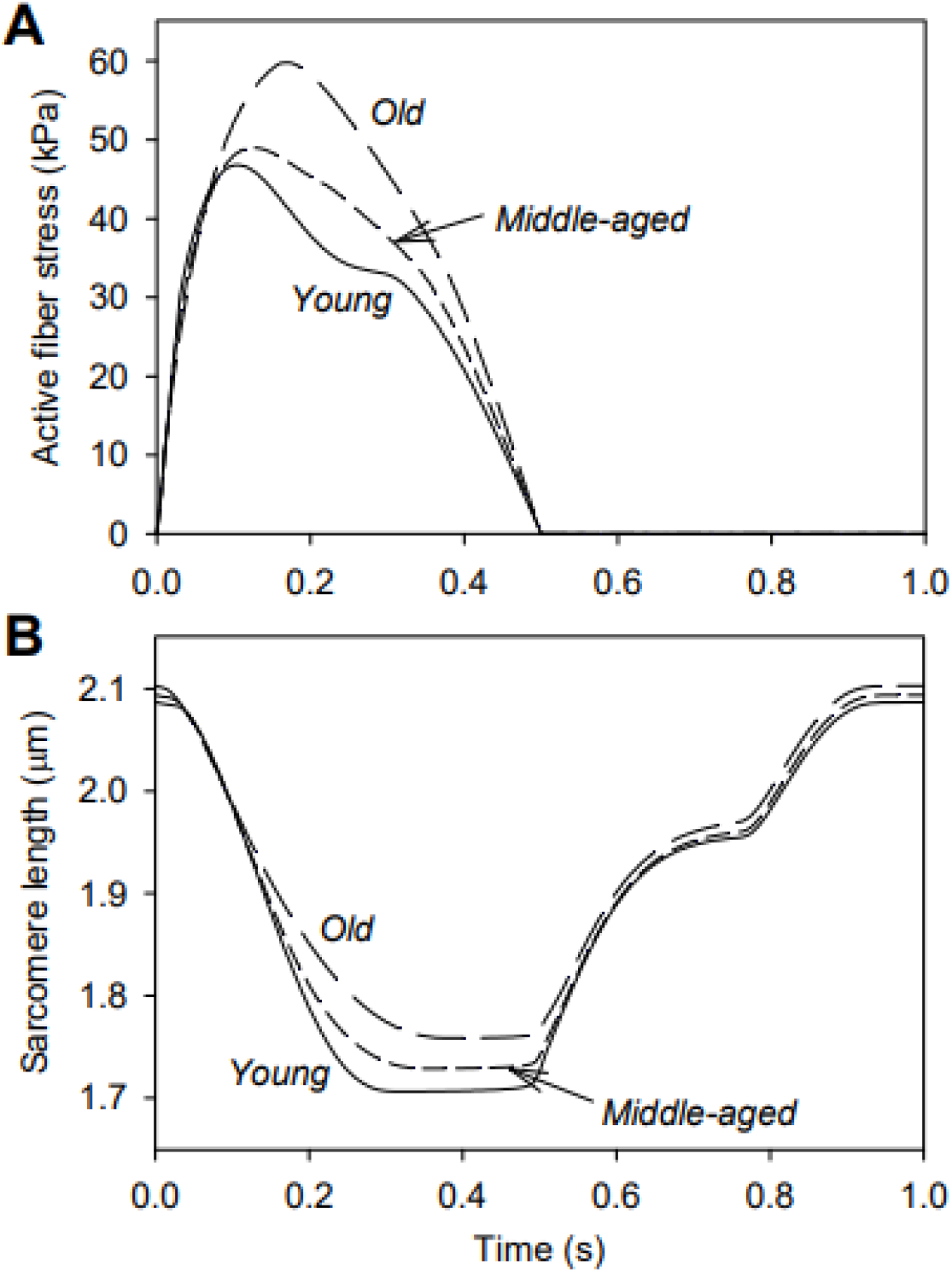
Effects of aorta aging on myocardial kinematics and dynamics. **A.** With aortic aging, the increase in aortic pressure during systole is balanced by increased active fiber stress, which is associated with slower shortening and increased sarcomere lengths. **B.** With aortic aging, the rate of sarcomere shortening during systole is decreased, with increased sarcomere lengths over most of the cycle. Sarcomere recovery after contraction is similar in all three aortic conditions.

### 3.4 Effects of reduced LV contractility combined with aging aorta

To examine effects of reduced LV contractility, three levels of maximal force generation are considered: *normal* (*k_mLV_* = 200 kPa), *impaired* (125 kPa) and *failing* (50 kPa). In each case, *k_vLV_* is altered in the same proportion, to maintain an appropriate force-velocity relationship. Figure 6 shows the resulting PV loops for each level of aortic aging. The simulations with varying aortic impedance allow estimation of the LV end-systolic pressure-volume relationship (ESPVR), defined as the slope of the line through the points of maximal elastance (P/V). The values are 2.4 (*normal*), 2.0 (*impaired*) and 0.88 (*failing*) in units of mmHg/cm^3^ and correlate with the assumed levels of contractility, as expected.

**Figure 6.**
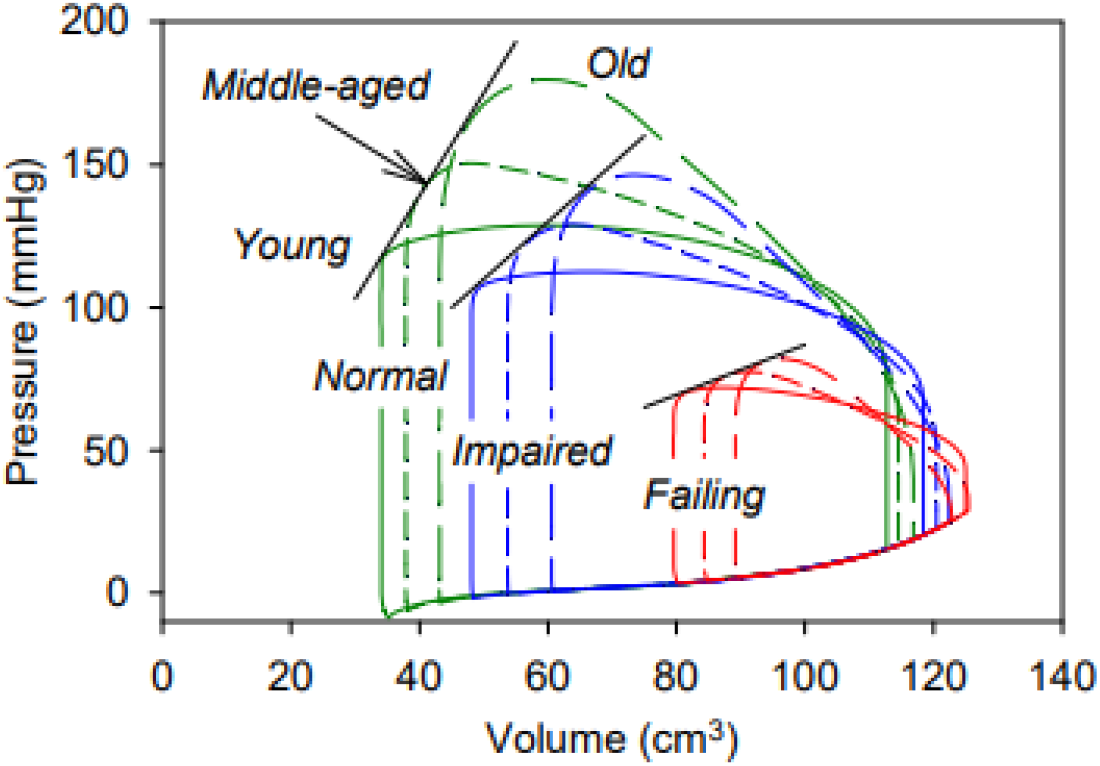
Combined effects of aortic aging and decreased LV contractility on PV loops. For each level of contractility, the end-systolic pressure-volume relationship is represented by a straight line.

The effects of reduced contractility and aging aorta on stroke volume are summarized in Figure 7A. With *impaired* contractility, the loss of contractile force is somewhat compensated by a shift to higher sarcomere lengths, i.e. up the length-tension curve, as a result of increased LV volume (Figure 6) and the reduction in stroke volumes is only 10-15%. With *failing* contractility, this mechanism cannot compensate effectively, and stroke volumes drop to 40-50% of the normal values. Corresponding changes in peak aortic pressure are shown in Figure 7B. Again, aortic pressure is somewhat maintained with *impaired* contractility, but greatly reduced with *failing* contractility. Figures 7 shows that the effect of aging aortic properties is to maintain pressure in the face of reduced contractility, but at the expense of reduced stroke volume.

**Figure 7.**
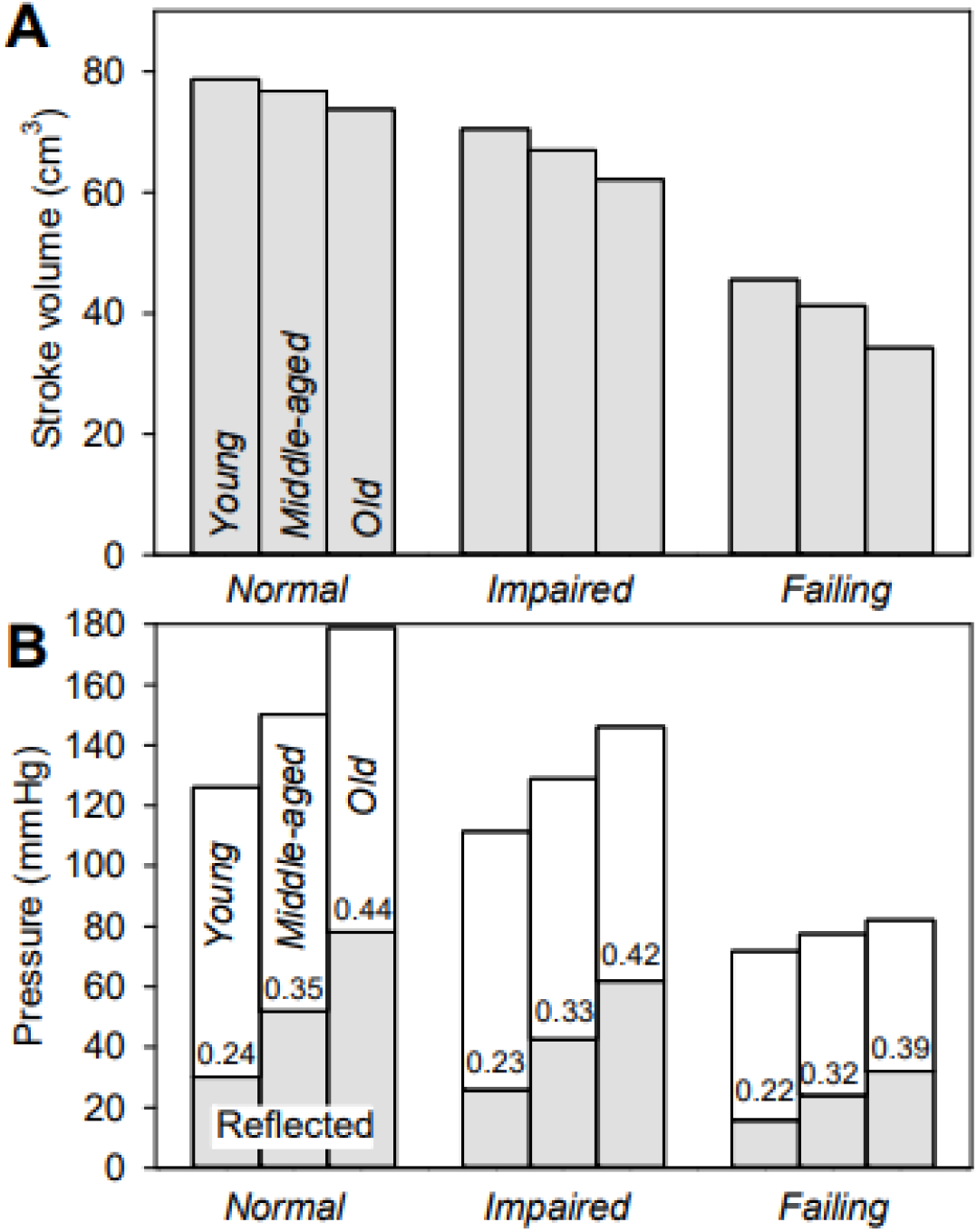
Combined effects of aortic aging and decreased LV contractility on stroke volume and maximum generated pressure. **A.** Stroke volume decreases with aortic aging. This effect becomes more significant with reduced LV contractility. **B.** Maximum generated pressure increases with aotic aging. This effect decreases markedly when LV contractility is reduced. Gray bars represent the component of pressure resulting from the reflected pressure wave, and numbers on each bar give the reflected component as a fraction of the total. In all cases, the reflected wave becomes more significant with aortic aging.

## 4. DISCUSSION

### 4.1 Model rationale

The circulatory system consists of multiple components, including the cardiac chambers and the various segments of the vasculature, whose interactions are not easily understood intuitively or by qualitative arguments. Theoretical models are therefore widely used to gain insights into circulatory system functions. Such models generally fall into two broad categories. Firstly, lumped or zero-dimensional models treat the circulatory system as an ensemble of relatively simple interconnected elements, such as a varying elastance model for the LV or a Windkessel model for the aorta. These models are computationally tractable but the lumped-element properties may not be directly related to measurable biophysical properties. Secondly, spatially resolved models take into account the detailed structure and properties of the heart and vasculature, but cannot represent the entire system at the same level of detail. Such models generally have high computational costs.

The present approach represents a middle path between these approaches. Critical components of the system, including the LV and aorta, are represented by spatially resolved models and coupled to lumped-element models for other system components. The spatially resolved models are designed to incorporate underlying biophysical properties, while allowing fast computation. With this approach, rapid scans over multiple sets of parameter values are facilitated. An eventual goal of this work is to estimate underlying cardiac and vascular properties online, using parameter estimation based on clinically available measurements, which can only be achieved if each model run takes a minimal computational time.

Relative to our previously published work (Moulton et al. 2017; Hong et al. 2019), the present model includes several significant new components: (i) 1-D wave propagation model for aorta; (ii) volume-preserving thick-walled spherical chamber models for the LA, RA and RV; (iii) zero-dimensional Bernoulli-type valve models; (iv) closed-loop circulation using lumped elements.

### 4.2 Left ventricle-aorta coupling

The model presented here is suitable for simulating ventricular-vascular interactions, because both the LV and the aorta are represented by biophysically based, spatially resolved models. The model includes two-way interaction: the flow in the aorta is driven by the LV, and the flow generated by the LV is affected by the reflected pressure wave in the aorta. In the model for the aorta, the reflected wave is assumed to be generated continuously along the length of the aorta as a result of its taper, which is represented by a decaying exponential dependence of cross-section area on distance. While arterial branch points also contribute to reflected waves, these effects are not explicitly represented in the model.

The main contribution of the aorta to the afterload on the LV is the reflected pressure wave. In the present model, this is computed by convolution of the impulse response function *f_refl_*(*t*) with the aortic inflow rate. As shown in Figure 4A, the response function has a complicated shape, resulting from wave propagation and reflection along the length of the aorta. The peak value is reached at a time 2*L / c*, where *c* is the wave speed. The shape of the function depends on the cross-section area and compliance of the aorta. If a Windkessel model were used instead of the spatially resolved model, these graphs would show mono-exponential decay. In comparison, the computed functions show significant effects of wave reflection. Figure 4B shows the important contribution of the reflected pressure pulse to the overall aortic pressure waveform.

### 4.3 Aging aorta

With aging, the aorta shows a marked increase in stiffness, and also increases in diameter, tortuosity, length, and taper (Hickson et al. 2010). Pulse wave velocity (PWV) increases by more than two-fold in the aging aorta because of the increase in stiffness (Phan et al. 2016; Hickson et al. 2010). This trend is only partially counteracted by the increase in diameter. The overall effect of aging is the generation of substantially larger wave reflections, which arrive earlier in systole (Figure 4) (Pagoulatou and Stergiopulos 2017; Heusinkveld et al. 2019; Safar 2008).

The changes in aortic properties have several effects on the function of the LV and circulatory system (Park et al. 2020). With aging, the increased impedance of the aorta leads to increased peak pressure in late systole, but decreased stroke volume (Figure 3). Pulse pressure increases substantially (Figure 4B), as is typically seen in aging patients (Sweitzer et al. 2013). As a result of the more rapid decline of pressure in diastole, the aortic valve opens earlier, giving an earlier upswing in aortic pressure in *old* versus *young* aorta (Figure 4B). The aortic flow waveform in systole is blunted but lengthened (Figure 4C, D).

At the level of the sarcomeres, the changes in aortic properties with aging have significant effects. The rate of contraction is reduced due to the increased afterload, and force generation is thereby increased according to the force-velocity relationship (Figure 5). In a recent study using extensively instrumented dogs, Park et al. (Park et al. 2020) found that early reflected waves generated by aortic occlusion slowed minor axis LV shortening, in agreement with the present predictions.

### 4.4 Interaction of failing heart and aging aorta

As shown in Figure 7A, the effects of aging aorta on stroke volume are relatively mild (6% decrease) with *normal* LV contractility, but become more severe (25% decrease) if contractility is *failing*. In the latter case, the LV has reached the limit of expansion in response to increased preload and cannot compensate for the reduction in ejection resulting from increased afterload. Maximum aortic pressure increases with aging aorta with *normal* and *impaired* LV contractility, but this effect is blunted when contractility is *failing*. This prediction is consistent with results from a clinical study of patients with systolic dysfunction (LVSD) (Paglia et al. 2014), where it was found that increased arterial wave reflections measured by increased central pulse pressure reduced LV flow in the patients without a positive impact on blood pressure. The *failing* LV is functioning at a higher part of the passive length-tension curve (Figure 6), and cannot produce more force by further extending to a higher point on the active length-tension curve (Figure 2).

### 4.5 Limitations

As in all models, this model includes multiple simplifications. The LV is assumed axisymmetric, and changes in regional function are not explored. The present approach allows inclusion of 3-D deformation modes (Hong et al. 2019). Effects of direct mechanical interactions between left and right heart are not included, but can be significant. The effects of aortic branching are not explicitly included, and all reflections are attributed to the taper of the aorta. A more comprehensive aortic model that includes a vascular tree (Heusinkveld et al. 2019) would allow analysis of the effects of aortic branching and would allow correlation with clinical measurements of pulse transit time.

### 4.6 Conclusion

A novel circulatory system model is presented, using a combination of spatially resolved and lumped-parameter component representations, which allows simulations of multiple cardiac cycles with computations that run faster than real time. The model is applied to examine the consequences of the aortic stiffening that occurs with aging, which increases aortic impedance and hence the afterload on the LV. The results emphasize the contribution of the reflected pressure wave to LV afterload, and show that this reflected wave increases in amplitude with aortic aging, causing increases in sarcomere length and fiber stress in the LV. The resulting deficit in stroke volume is relatively small in the normal heart, but is amplified if LV contractile force is low in heart failure. Because of its fast computational speed and comprehensive representation of system phenomena, this model has potential applications for exploring other aspects of circulatory system performance, and to online estimation of underlying parameters from clinically available measurements.

## Compliance with ethical standards

### Funding

Timothy W. Secomb is supported by NIH Grant U01 HL133362.

### Animal Studies

This study does not contain studies with animals by any of the authors.

### Human subjects

The study protocol was approved by the Institutional Review Board of the University of Nebraska Medical Center. All procedures followed were in accordance with the committee on human experimentation (institutional and national) and with the Helsinki declaration of 1975, as revised in 2000. Written informed consent was obtained from the volunteer before the study.

### Author Contribution Statement

M.M. and T.S. developed the LV mathematical model. T.S. developed the aorta pulse wave model. M.M. and T.S. wrote the manuscript, edited the manuscript and prepared the figures. All authors reviewed the final manuscript.

