## Supplementary Material for "A fast computational model for circulatory dynamics: Effects of left ventricle-aorta coupling"

Michael J. Moulton and Timothy W. Secomb

**A. MODEL FOR LV KINEMATICS AND DYNAMICS**

**Deformation gradient tensor and Green strain tensor.** Prolate spheroidal coordinates (*μ*,*ν*,*φ*) are used with a time-dependent interfocal distance 2*a*. Then
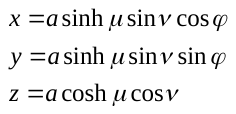
(S1)

where (*x*,*y*,*z*) are Cartesian coordinates. The scale factors are
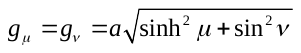
 and
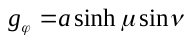
, and the Jacobian is

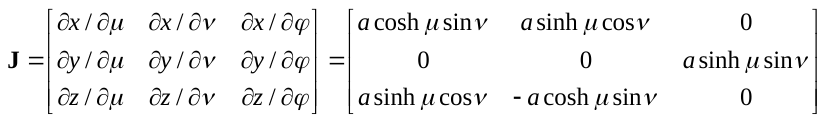
(S2)

where *φ* = 0 without loss of generality**.** The determinant is

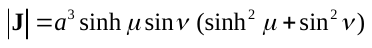
(S3)

A reference configuration is defined with *a* = *a*_0_ and coordinates (*μ*_0_,*ν* _0_,*φ* _0_), in which the LV wall occupies the region
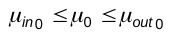
 and
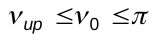
. The deformation is represented using a family of mappings (*a*_0_;*μ*_0_,*ν* _0_,*φ* _0_) → (*a*;*μ*,*ν* ,*φ* ). For volume conservation,

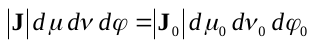
(S4)

where **J**_0_ is the Jacobian in the reference configuration. The assumption of axisymmetry implies that
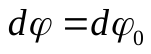
 and we further assume that
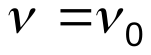
. Then (S3) and (S4) imply that

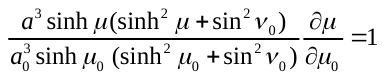
(S4.1)

yielding an implicit relation between *μ* and *μ* _0_:

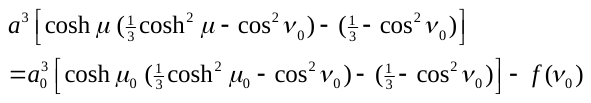
(S4.2)

where
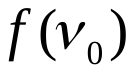
 is an arbitrary function. A parameter *a*_1_ where *a* = *a*_0_ + *a*_1_ is introduced to represent ventricular lengthening. The function
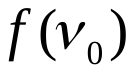
 is chosen as a function of a parameter *a*_2_ so that the internal volume approaches zero as *a*_2_ → *a*_0_, corresponding to 100% ejection. The relation between
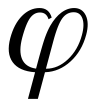
 and
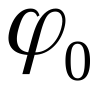
 is chosen to represent ventricular torsion dependent on a parameter *a*_3_. The mapping is thus defined by:

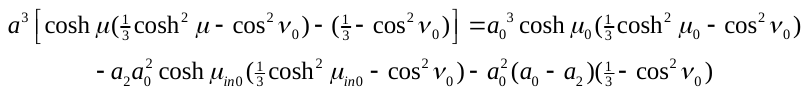
(S5)

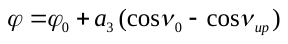
 (S6)

. The deformation gradient tensor in prolate spheroidal coordinates is:

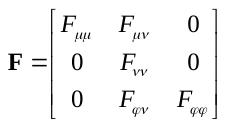
 (S7)

The components are

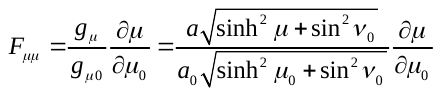
(S8)

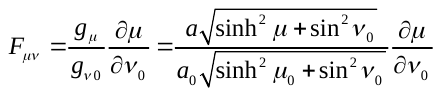
(S9)

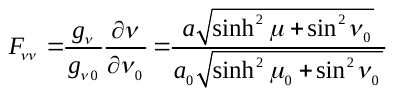
(S10)

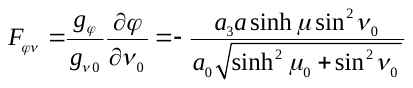
(S11)

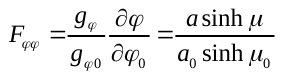
(S12)

To evaluate these components and their derivatives, the following results are needed:

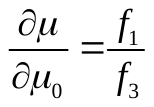
 and
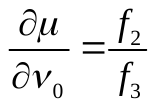
(S13)

where

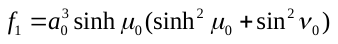
(S14)

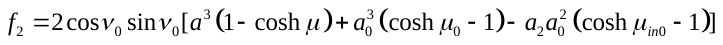
(S15)
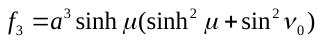
(S16)

It follows that

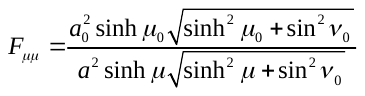
(S17)

and
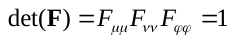
 as expected. The right Cauchy-Green tensor is
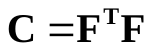
. The components of the Green strain tensor
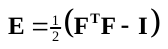
 are:

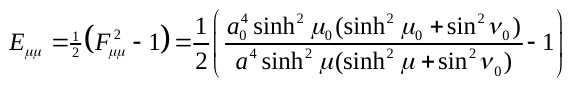
(S18)

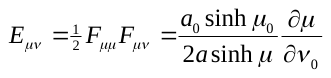
(S19)

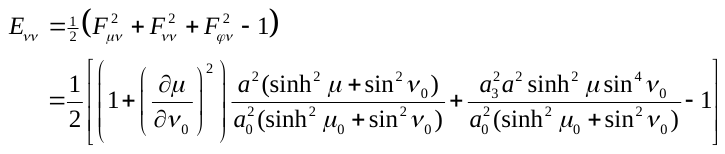
(S20)

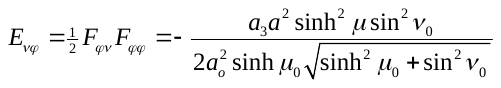
(S21)

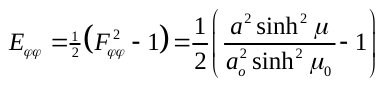
(S22)

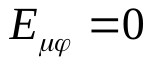
(S23)

**Derivatives of the Green strain tensor.** The derivatives with respect to *a*_1_, *a*_2_ and *a*_3_ are calculated from the derivatives of the deformation gradient tensor. The following results are needed:

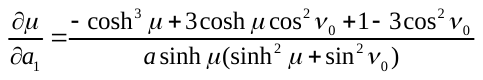
(S24)

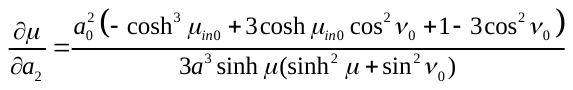
(S25)

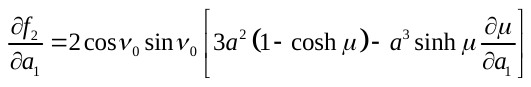
(S26)

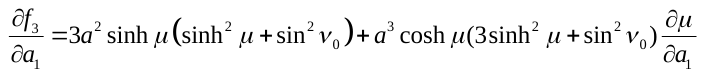
(S27)

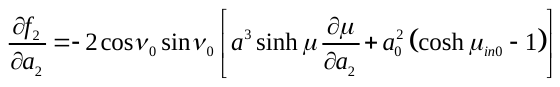
 (S28)

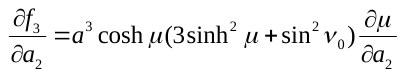
(S29)

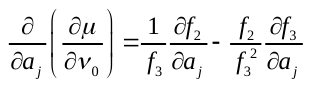
 and
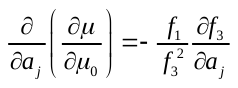
 for
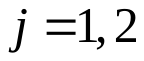
(S30)

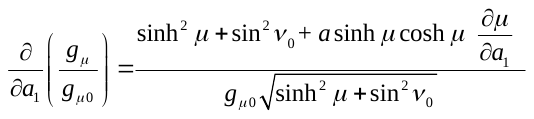
(S31)

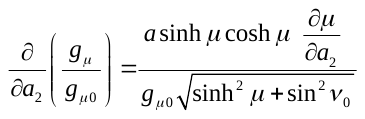
(S32)

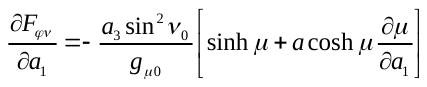
,
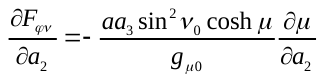
(S33)

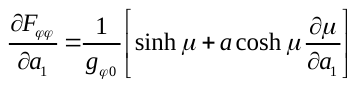
,
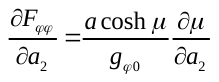
(S34)

The derivatives of components of **E** are deduced from the derivatives of **F**.

**Base vectors in prolate spheroidal and fiber coordinate systems.** In the reference configuration, the base vectors
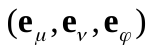
 of the prolate spheroidal coordinate system (where *φ* = 0 without loss of generality) are:

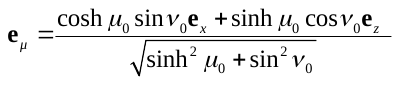
(S35)
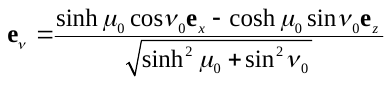
(S36)

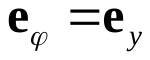
(S37)

In the reference configuration, muscle fibers are assumed to lie in surfaces of constant
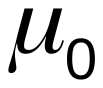
, and are identified by
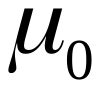
 and by their angular position
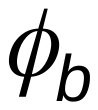
 at the base of the heart. Position on a given fiber is parameterized by
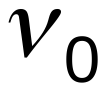
. The helical arrangement of the fibers and the variation of the helix angle through the wall are represented by setting
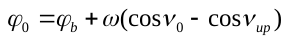
, where
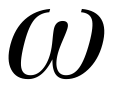
 is a
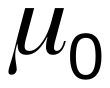
-dependent wrapping parameter. The fiber paths in the reference configuration are expressed parametrically by

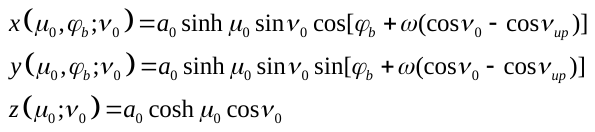
(S38)

A local coordinate system
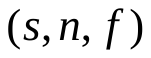
 is defined based on the reference fiber geometry, where
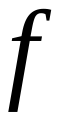
is the fiber direction,
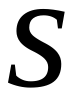
is the transmural direction and
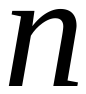
is perpendicular to
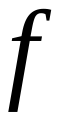
 and
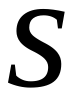
. The base vectors
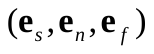
 of the fiber coordinate system (where *φ* = 0) are:

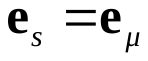
(S39)

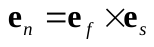
(S40)

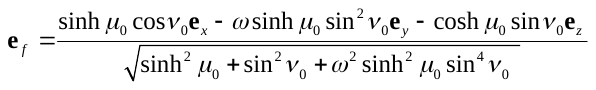
(S41)

The angle
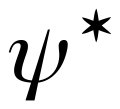
 is defined as the angle that the negative fiber direction
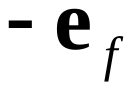
 makes with the
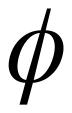
-coordinate direction
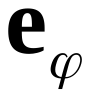
 in the reference configuration. Therefore

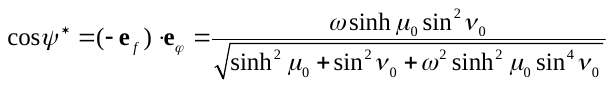
(S42)

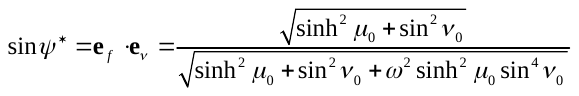
(S43)

The orientation of the fibers relative to the
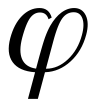
-coordinate direction is measured either by
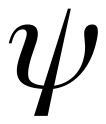

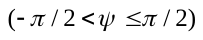
 or by
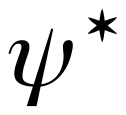

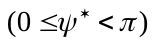
, where
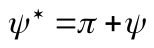
 if
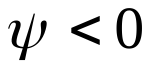
 and
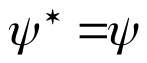
if
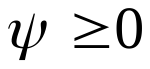
. The fiber angle
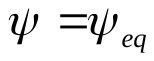
 at the equator of the LV
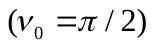
 varies from
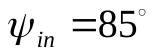
 on the inner wall to
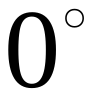
 at the midwall and to
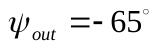
 on the outer wall. This is represented assuming linear variation with
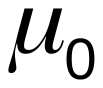
, according to

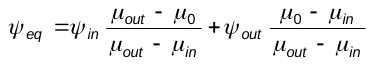
 (S24.1)

When
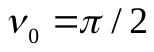
, (S23) gives

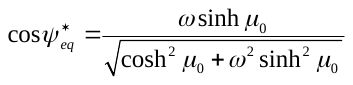
(S24.2)

which can be solved for the wrapping parameter,

if	(S24.3)

Using the definition of the base vectors, a rotation matrix is derived to convert vectors and tensors from the fiber Cartesian system to the prolate direction:

(S25)

To compute the work integrals (see below), the components of the stress tensors are converted from fiber coordinates to the undeformed prolate coordinates . The components of a PK2 stress in prolate coordinates are related to the components in fiber coordinates by

**Virtual work on the boundary.** The load on the boundary consists of the internal pressure . The virtual work can therefore be calculated as the pressure multiplied by the change of volume. The ventricular volume is computed as a solid of revolution:

(S56)

The volume change is where the integrals are evaluated on :

(S57)

(S58)

**Passive stress.** The passive myocardium is assumed to be viscoelastic, with the total stress as the sum of elastic and viscous components. The elastic stress is given by a strain-energy function

(S59)

where	(S60)

are the Green strain components in the fiber coordinates and are material parameters. The PK2 stress is

(S61)

The viscous stress is calculated using the equations for a viscous fluid:

(S62)

where in Cartesian coordinates. Then letting

(S63)

gives	.	(S64)

The PK2 stress satisfies

(S65)

where is the finger strain and and . Therefore,

(S66)

In this case, *J* = 1. Since is symmetric, this can be written

.	(S67)

This can be expressed in terms of the derivatives of :

(S68)

where	(S69)

**Active fiber stress.** The Cauchy stress generated by active fiber contraction is approximated by:

(S70)

Here, gives the maximum active stress, is the fiber strain, defines the force-velocity characteristics of the fibers and determines the sensitivity of active force to preload according to the Frank-Starling mechanism, where is the end-diastolic fiber strain. The function

(S71)

describes the length-tension characteristics of active force generation, where is sarcomere length, is the length at which maximum tension is generated and determines the width of the peak in the curve. The time-dependent fiber activation is given by

(S72)

where is time after the start of contraction in a given cycle, is the period of activation and is the period of the cardiac cycle. The exponent steepens the rise and fall of activation with increasing end-diastolic strain . The resulting Cauchy stress is where denotes the tensor product and is the basis vector for the local fiber direction. The corresponding PK2 stress is , which can be written:

(S73)

where	(S74)
for	(S75)

**Force balance equations.** The equations of mechanical equilibrium in weak form imply that, for each of the parameters *a_j_*, the variation in internal work with respect *a_j_* must equal the rate at which work is done by external traction, which in this case is the pressure *P_lv_*, i.e.

for	(S76)

Because the stress has viscous and elastic components, this can be restated as a differential equation for *a_i_* where the coefficients are the incremental virtual work integrals. The virtual work integrals are given in terms of the PK2 stresses and Green strains as

(S77)

With these definitions, the equations for the LV can be written as a system of three differential equations

(S78)

**B. MODEL FOR LA, RV AND RA KINEMATICS AND DYNAMICS**

**Kinematics.** The left and right atria are modeled as spherical shells, and the right ventricle is modeled as a segment of a spherical shell. In a contracting sphere, if the point *r*_0_ is mapped to *r*, conservation of volume implies that and so where are constants of integration, with for the LA, RV, RA respectively. The stretch ratio in the surface is , the deformation gradient tensor is and the Green’s strain tensor is

.	(S79)

Note that , so

.	(S80)

**Passive stress.** The passive elastic strain energy density is where . The PK2 stress is

.	(S81)

The passive viscous stress is where , and , yielding

where .	(S82)

**Active stress.** The active Cauchy stress is given by where and is the fiber strain. The factor ½ is introduced because the stress is assumed to be isotropic in the surface instead of unidirectional. Note that . The corresponding PK2 stress can be expressed as , where

(S83)

(S84)

The coefficients in the differential equations are

,,	(S85)

The equations governing the internal dynamics of the LA, RV and RA are of the form:	,	(S86)

where is the pressure in the chamber.

**C. 1-D MODEL FOR AORTA**

A cylindrical aorta is assumed with cross-section area *A*(*x*,*t*), flow rate *q*(*x*,*t*), and pressure *p*(*x*,*t*), where *x* is position and *t* is time. Blood velocity is approximated by flow rate divided by area, and advective acceleration is neglected. Conservation of fluid mass and momentum imply that

(S87)

(S88)

where *ρ* is density. The compliance of the aorta is defined as

(S89)

and is assumed constant. A reference state is defined with zero flow, and a cross-section area that varies with position, , where *A*_00_ is the area at *x* = 0 and *A*_01_ gives the degree of taper. Equations (S87) and (S88) are linearized about this state, giving

(S90)

(S91)

The wave speed is and the wave impedance is .

For the numerical analysis, pressures (*p_i_*) are computed at *n* nodes, evenly distributed with spacing *∆x* along the length of the aorta, and flows (*q_i_*) are computed at *n*−1 nodes lying midway between the pressure nodes. Equations (S90) and (S91) are represented using centered differences as

(S92)

(S93)

At the inlet *x* = 0, a flow boundary condition is imposed, . Equation (S93) then gives

(S94)

and, using equation (S92),

(S95)

At *x* = *L*, a matched impedance boundary condition is applied, to avoid a discrete reflection:

(S96)

To analyze the impulse response, the system (S92), (S93), (S95) and (S96) is solved for *p_i_*(*t*) and *q_i_*(*t*), assuming a short pulse of flow with unit weight:

(S97)

where *t_p_* << *t_c_*, the cardiac period. Fourth-order Runge-Kutta integration is used with time step ∆*t*_1_. Typical values are *t_p_* = 0.02 s, *L* = 50 cm, *n* = 501, ∆*x* = 0.1 cm and ∆*t*_1_ *=* 10^-4^ s. A compliance *G*_0_ = 6.25 × 10^−6^ (dyn/cm^2^)^−1^ with *ρ* = 1.05 g/cm^3^ yields a wave speed c = 390 cm/s. The integration is continued for a time 3*t_c_*, which is much larger than *L*/*c*, the time for the pulse wave to traverse the aorta, so that reflected waves after 3*t_c_* are negligible.

Two impulse-response functions describe the behavior of the aorta in this model: the reflected pressure *f_refl_*(*t*) at *x* = 0 and the transmitted flow *f_trans_*(*t*) at *x* = *L*. Because the input flow is periodic, the responses during the second and third cardiac periods are added to those in the first period, giving the response over a single cycle to a periodic input. To obtain *f_refl_*, the pressure spike *Z*(0)*q_in_* resulting from the imposed flow pulse is subtracted from *p*_1_(*t*). The time-average of *f_refl_* is then subtracted, so that the resulting function has zero mean. The functions *f_refl_* and *f_trans_* depend only on the properties of the aorta and are calculated prior to the simulation of the circulatory system. Their values are saved at time steps of the main model simulation.

Given these impulse-response functions, the response of the aorta to an arbitrary inflow waveform *q_lv_*(*t*) can be computed by convolution. The reflected pressure and the transmitted flow are

(S98)	(S99)

where *q_lv_* is the flow rate at *x* = 0. In this system, the pressure is arbitrary up to an additive constant. To remove this arbitrariness, the impedance of the aorta to the steady component of flow is assumed to be negligible, implying that where the overbar denotes the average over one cardiac cycle. The pressure at *x* = 0 can then be computed as

(S100)

The quantities and represent averages over one cycle period of the corresponding time-dependent variables, and are continuously updated as the calculation proceeds.

**D. MODEL FOR VALVES**

The four cardiac valves are represented by a zero-dimensional (0-D) Bernoulli-type model. Flows exiting the LV, LA, RV and RA are governed respectively by

(S101)

where the inertial coefficients are

(S102)

and the kinetic energy coefficients are

(S103)

The valve areas vary with time and are given by

(S104)

where the *ζ* variables are governed by

(S105)

(S106)

(S107)

(S108)

**E. MODEL INCLUDING SYSTEMIC AND PULMONARY CIRCULATIONS**

The overall model configuration is shown in Figure 1. Variables are defined such that resistances and flow rates are downstream from the corresponding nodal pressures. The equations of mechanical equilibrium for the four chambers of the heart are:

(S109)

(S110)

(S111)

Conservation of volume for the four heart chambers gives:

(S112)

(S113)

(S114)

(S115)

The factor *f_rv_* gives the fraction of the sphere representing the RV. Conservation of volume for the four compliances representing the systemic arteries, systemic veins, pulmonary arteries and pulmonary veins gives:

(S116)

(S117)

(S118)

(S119)

The inertia of the systemic and pulmonary veins is included:

(S120)

(S121)

Equations (S101), (S105-111) and (S113-121) define the derivatives of 20 dynamical variables:

(S122)

The values of are obtained by solving the system of four linear algebraic equations (S109)-(S112). From (S86), the other pressures in the system are given by:

(S123)

(S124)

(S125)

(S126)

and equation (S100) for . The resulting system of 20 ordinary differential equations is integrated using a standard RK2 scheme with a time step s.
